# Omicron BA.1 and BA.2 neutralizing activity elicited by a comprehensive panel of human vaccines

**DOI:** 10.1101/2022.03.15.484542

**Authors:** John E. Bowen, Kaitlin R. Sprouse, Alexandra C. Walls, Ignacio G. Mazzitelli, Jennifer K. Logue, Nicholas M. Franko, Kumail Ahmed, Asefa Shariq, Elisabetta Cameroni, Andrea Gori, Alessandra Bandera, Christine M. Posavad, Jennifer M. Dan, Zeli Zhang, Daniela Weiskopf, Alessandro Sette, Shane Crotty, Najeeha Talat Iqbal, Davide Corti, Jorge Geffner, Renata Grifantini, Helen Y. Chu, David Veesler

**Affiliations:** Department of Biochemistry, University of Washington, Seattle, WA 98195, USA; Howard Hughes Medical Institute, University of Washington, Seattle, WA 98195, USA; Instituto de Investigaciones Biomédicas en Retrovirus y SIDA (INBIRS), Facultad de Medicina, Buenos Aires C1121ABG, Argentina; Division of Allergy and Infectious Diseases, University of Washington, Seattle, WA 98195, USA; Department of Paediatrics and Child Health, and Biological & Biomedical Sciences, Aga Khan University, Karachi 74800, Pakistan; Humabs Biomed SA, a subsidiary of Vir Biotechnology, 6500 Bellinzona, Switzerland; Infectious Diseases Unit, Foundation IRCCS Ca’ Granda Ospedale Maggiore Policlinico, Milan, Italy; Department of Pathophysiology and Transplantation,, University of Milan, Milan, Italy; Centre for Multidisciplinary Research in Health Science (MACH), University of Milan, Milan, Italy; Vaccine and Infectious Disease Division, Fred Hutchinson Cancer Research Center, Seattle, WA, USA; Center for Infectious Disease and Vaccine Research, La Jolla Institute for Immunology, La Jolla, CA 92037, USA; Department of Medicine, Division of Infectious Diseases and Global Public Health, University of California, San Diego, La Jolla, CA UC92037, USA; INGM, Istituto Nazionale Genetica Molecolare “Romeo ed Enrica Invernizzi”, 20122 Milan, Italy

## Abstract

The SARS-CoV-2 Omicron variant of concern comprises three sublineages designated BA.1, BA.2, and BA.3, with BA.2 steadily replacing the globally dominant BA.1. We show that the large number of BA.1 and BA.2 spike mutations severely dampen plasma neutralizing activity elicited by infection or seven clinical vaccines, with cross-neutralization of BA.2 being consistently more potent than that of BA.1, independent of the vaccine platform and number of doses. Although mRNA vaccines induced the greatest magnitude of Omicron BA.1 and BA.2 plasma neutralizing activity, administration of a booster based on the Wuhan-Hu-1 spike sequence markedly increased neutralizing antibody titers and breadth against BA.1 and BA.2 across all vaccines evaluated. Our data suggest that although BA.1 and BA.2 evade polyclonal neutralizing antibody responses, current vaccine boosting regimens may provide sufficient protection against Omicron-induced disease.

---

The ongoing COVID-19 pandemic has led to the emergence of SARS-CoV-2 variants with increased transmissibility, viral fitness and immune evasion (*1–10*). The most recently named variant of concern, Omicron, is characterized by the greatest known genetic divergence from the ancestral virus (Wuhan-Hu-1) and consists of three sublineages designated BA.1, BA.2, and BA.3. These three sublineages are as genetically and antigenically distant from each other as several previous SARS-CoV-2 variants of concern are from one another (*11*). BA.1 was first detected in late 2021 and quickly displaced Delta (*3, 9, 12*) to become the globally dominant SARS-CoV-2 strain, aided by its high transmissibility and escape from neutralizing antibodies (*6, 13–17*). However, BA.2 has recently been shown to replicate with faster kinetics, mediate enhanced fusogenicity, and be more pathogenic than BA.1 in hamsters (*18, 19*), in agreement with its steadily increasing prevalence and expected future global replacement of BA.1 (*20*).

The receptor-binding domain (RBD) of the SARS-CoV-2 spike (S) glycoprotein interacts with the receptor ACE2 (*21–25*), promoting S conformational changes that lead to membrane fusion and viral entry (*26, 27*). S is the main target of neutralizing antibodies, which have been shown to be a correlate of protection against SARS-CoV-2 (*28–36*), with RBD-targeting antibodies accounting for nearly all cross-variant neutralizing activity (*37*). SARS-CoV-2 vaccines are thus based on the S glycoprotein (sometimes the RBD only (*28, 38, 39*)) or full virus (inactivated) and utilize a variety of delivery technologies. Lipid-encapsulated prefusion-stabilized S-encoding mRNA vaccines include Moderna mRNA-1273 and Pfizer/BioNTech BNT162b2. Viral-vectored vaccines encoding for the SARS-CoV-2 S sequence include Janssen Ad26.COV2.S (human adenovirus 26), AstraZeneca AZD1222 (chimpanzee adenovirus), and Gamaleya National Center of Epidemiology and Microbiology Sputnik V (human adenovirus 26/5 for prime/boost). Novavax NVX-CoV2373 is a prefusion-stabilized S protein-subunit vaccine formulated with a saponin-based matrix M adjuvant whereas Sinopharm BBIBP-CorV comprises inactivated virions. The primary vaccine series consisted of two doses for all of these candidates except for Ad26.COV2.S which was administered as a single dose.

We set out to compare side-by-side the plasma neutralizing activity elicited in humans by each of these seven vaccines or SARS-CoV-2 infection and evaluate the immune evasion associated with the constellation of spike mutations present in the BA.1 and BA.2 Omicron sublineages. Although BA.1 and BA.2 share a large number of spike mutations, they are each characterized by unique sets of amino acid changes, which are expected to be associated with different antigenic properties (**Table S1**). We therefore assessed entry of vesicular stomatitis virus (VSV) pseudotyped with the SARS-CoV-2 Wuhan-Hu-1 S harboring the D614G, BA.1, or BA.2 mutations into VeroE6 cells stably expressing TMPRSS2 (*40*) in the presence of vaccinee or convalescent plasma. Convalescent plasma was obtained from individuals infected with a Washington-1-like SARS-CoV-2 strain based on time of infection. Although we determined a plasma neutralizing geometric mean titer (GMT) of 162 against G614 VSV S pseudovirus, only 6/14 and 7/14 individuals had detectable, but mostly weak, neutralizing activity against BA.1 and BA.2, respectively. Individuals who received two doses of mRNA vaccines three to four weeks apart, however, fared better against Omicron sublineages than these previously infected patients. Subjects that received two doses of Moderna mRNA-1273 had G614, BA.1, and BA.2 S VSV neutralizing GMTs of 1155, 26, and 47, respectively, whereas subjects that received two doses of Pfizer BNT162b2 had G614, BA.1, and BA.2 S VSV neutralizing GMTs of 764, 23, and 34, respectively. 18/21 and 19/21 mRNA-vaccinated subjects retained neutralizing activity against BA.1 S VSV and BA.2 S VSV, respectively, with the combined cohorts experiencing a ≥35-fold GMT reduction against BA.1 S VSV and a ≥24-fold GMT reduction against BA.2 S VSV. A similar trend was observed for Novavax NVX-CoV2373 vaccinees (two doses) for which we determined a neutralizing GMT of 289 against G614 S VSV with only 1/13 and 5/13 individuals having detectable neutralizing activity against BA.1 (GMT : 11, ≥23-fold drop) and BA.2 (GMT : 18, ≥13-fold drop).A single dose of Janssen Ad26.COV2.S resulted in a G614 S VSV GMT of 80 and only 1/10 subjects had detectable plasma neutralizing activity against either Omicron sublineage. Two doses of AZD1222 four weeks apart induced G614, BA.1 and BA.2 S VSV neutralizing GMTs of 656, 15, and 31, amounting to ≥45-fold and ≥21-fold magnitude reductions for BA.1 and BA.2, respectively. Sputnik V vaccinee plasma after two doses had a G614 S VSV GMT of 153 and detectable neutralizing activity for 7/12 subjects against the two Omicron sublingeages, with ≥12-fold BA.1 and 7-fold BA.2 GMT respective reductions. Lastly, plasma from subjects vaccinated with two doses of Sinopharm BBIBP-CorV had a neutralizing GMT against G614 S VSV of 188 with only 5/12 and 8/12 samples retaining detectable neutralizing activity against BA.1 (GMT :15, ≥12-fold GMT reduction) and BA.2 (GMT:29, ≥6-fold GMT reduction), respectively. Overall, these data underscore the unprecedented magnitude of evasion of polyclonal plasma neutralizing antibody responses for these two Omicron sublineages in humans based on primary immunization regimes for all seven vaccines or infection with a more marked effect for BA.1 compared to BA.2. respectively. Overall, these data underscore the unprecedented magnitude of evasion of polyclonal plasma neutralizing antibody responses for these two Omicron sublineages in humans based on primary immunization regimes for all seven vaccines or infection with a more marked effect for BA.1 compared to BA.2.

**Figure 1.**
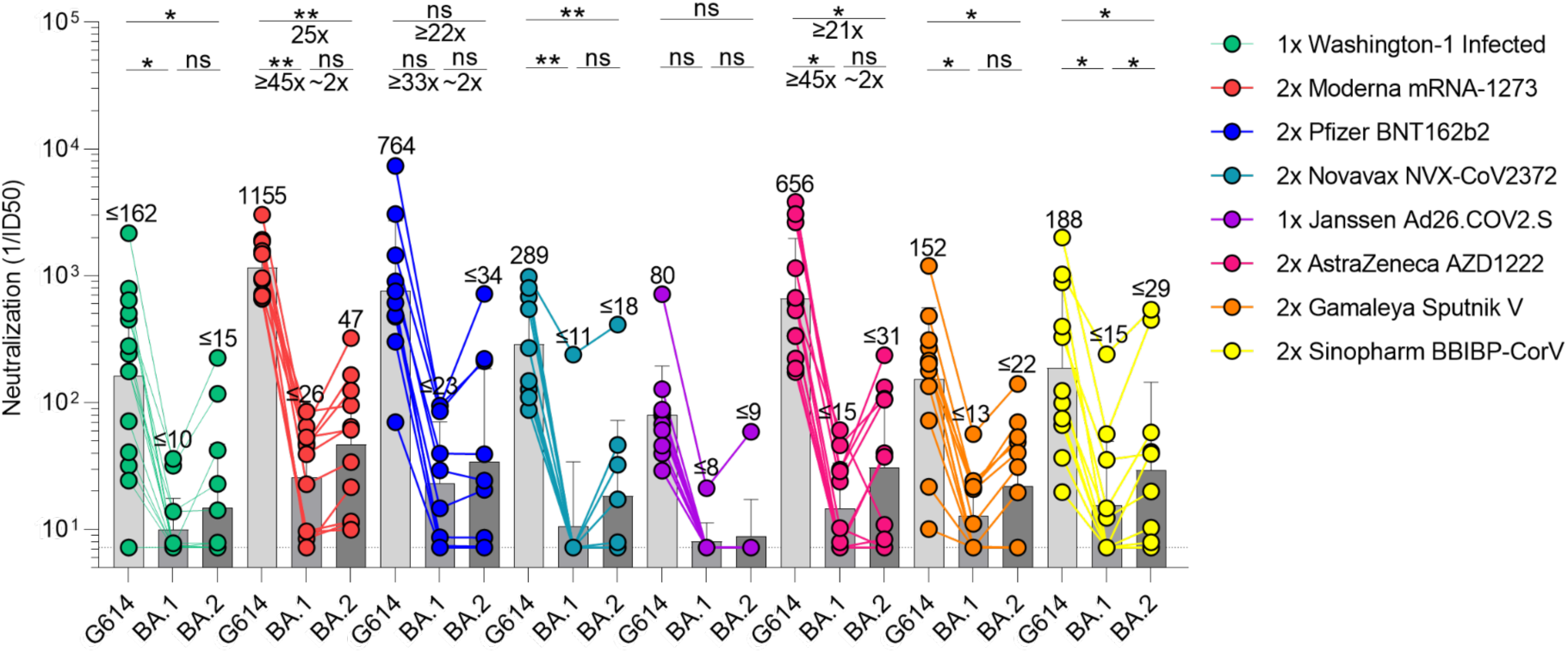
SARS-CoV-2 Omicron BA.1 and BA.2 evade human plasma neutralizing antibodies elicited by infection or primary vaccine series. Plasma neutralizing antibody titers elicited by primary COVID-19 vaccination determined using SARS-CoV-2 spike VSV pseudotypes using VeroE6-TMPRSS2 as target cells. Individual points are representative geometric mean titers from two independent experiments consisting of two replicates each. Bars represent geometric means and error bars represent geometric standard deviations for each group. Statistical significance between groups of paired data was determined by Wilcoxon rank test and *p< 0.05, **p< 0.01, ***p< 0.001, ****p< 0.0001. Patient demographics are shown in Table S2. Normalized curves and fits are shown in Figure S1.

The emergence of the SARS-CoV-2 Delta and subsequently Omicron variants of concern led to an increasing number of reinfections and vaccine breakthrough cases (*5, 41, 42*). Public health policies were therefore updated worldwide to recommend administration of an additional vaccine dose (booster) several months after the primary vaccine series, which has been shown to increase the breadth and potency of neutralizing antibodies (*5, 13*). We thus assessed and compared the benefits provided by homologous or heterologous vaccine boosters on vaccinee plasma neutralizing activity against G614, BA.1, and BA.2 S VSV pseudotypes. Plasma samples of subjects that received three mRNA vaccine doses had neutralizing GMTs of 3324, 415, and 612 against G614, BA.1, and BA.2 S VSV, respectively. The 8-fold and 5-fold respective potency losses against BA.1 and BA.2 are marked improvements over the corresponding ≥35-fold and ≥24-fold reductions observed after two vaccine doses, underscoring an increase in overall neutralizing antibody potency and breadth (*5, 13*). Plasma from individuals who received one dose of Ad26.COV2.S followed either by a homologous Ad26.COV2.S or a heterologous BNT162b2 booster approximately four months later had neutralizing GMTs of 641, 37 and 74 against G614, BA.1 and BA.2 S VSV, respectively, corresponding to dampening of ≥17-fold and 9-fold with 12/14 and 14/14 samples exhibiting detectable BA.1 and BA.2 neutralizing activity. We next investigated individuals that received two doses of AZD1222 four weeks apart followed by an mRNA booster approximately six months later. This cohort had neutralizing GMTs of 2579, 275, and 414 against G614, BA.1 and BA.2 S VSV, corresponding to 9-fold and 6-fold reductions against BA.1 and BA.2, respectively. These neutralizing GMT reductions are comparable to that observed with three doses of mRNA vaccines. Individuals vaccinated with two doses of Sputnik V and boosted with AZD1222 or BNT162b2 approximately nine months later had neutralizing GMTs of 1480, 160, and 147 for G614, BA.1, and BA.2, respectively, amounting to 9-fold and 10-fold reductions of potency against BA.1 and BA.2. The marked improvement in plasma neutralizing activity for subjects that received a booster dose over those that did not highlights the importance of vaccine boosters for eliciting potent neutralizing antibody responses against Omicron sublineages.

**Figure 2.**
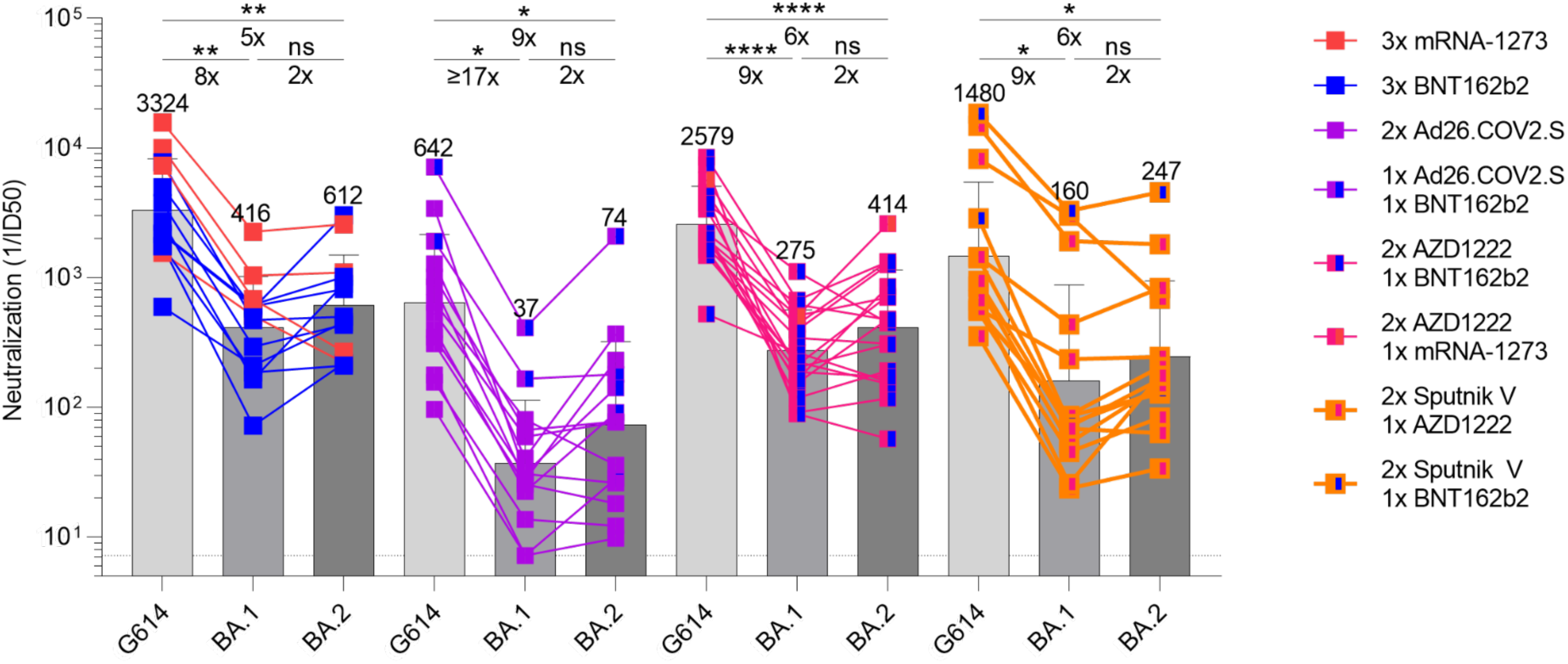
Booster doses rescue neutralization potency against Omicron BA.1 and BA.2. Plasma neutralizing antibody titers elicited by COVID-19 vaccine boosters determined using SARS-CoV-2 S VSV pseudotypes and VeroE6-TMPRSS2 as target cells. Individual points are representative geometric mean titers from two independent experiments consisting of two replicates each. Bars represent geometric means and error bars represent geometric standard deviations for each group. Statistical significance between groups of paired data was determined by Wilcoxon rank test and *p< 0.05, **p< 0.01, ***p< 0.001, ****p< 0.0001. Patient demographics are shown in Table S2. Normalized curves and fits are shown in Figure S2.

The development of life-saving vaccines is regarded as one of humanity’s greatest medical and scientific achievements, which is perhaps best exemplified by COVID-19 vaccines (*43–45*). Here we report that, although the two Omicron sublineages evaluated are characterized by severe dampening of plasma neutralizing activity, vaccine and infection-elicited cross-neutralization of BA.2 was more efficient than that of BA.1, independent of the immunization scheme or vaccine platform. Although mRNA vaccines induced the greatest magnitude of Omicron plasma neutralizing activity across all tested COVID-19 vaccines, administration of a booster dose increased neutralizing antibody titers and breadth against BA.1 and BA.2 to appreciable levels regardless of the vaccine evaluated, concurring with recent findings for BA.1 (*5, 13, 16, 46–48*). These results are consistent with previous studies demonstrating that a third vaccine dose results in the recall and expansion of pre-existing SARS-CoV-2 S-specific memory B cells, as well as de novo induction of novel ones, leading to production of neutralizing antibodies with enhanced potency and breadth against variants (*49, 50*). Furthermore, vaccinees receiving two doses of Ad26.COV2.S (four months apart) had greater Omicron neutralization potency than other two dose vaccine recipients (three to four weeks between doses) but less than three dose vaccinees. These findings suggest that the time interval between immunizations may affect the breadth and potency of elicited plasma neutralizing activity, and that a third dose may be beneficial for this cohort as well. Moreover, the induction by several currently available vaccines of robust cross-reactive cellular immunity against SARS-CoV-2 Omicron is likely playing a key role in the retained protection observed against severe disease (*51, 52*).

As SARS-CoV-2 becomes endemic in the human population, vaccination strategies will need to be carefully considered and optimized to provide long-lasting immunity. So far, elicitation of high titers of variant-neutralizing antibodies and protection against severe disease can be accomplished with three doses of the Wuhan-Hu-1 S antigen, as shown in animal models and studies of vaccine efficacy in humans(*53, 54*). In fact, an Omicron BA.1 S boost may not offer significantly more BA.1 protection than a Wuhan-Hu-1 S boost (*55–58*). However, continued SARS-CoV-2 evolution will accentuate the genetic drift from the ancestral strain and it is unknown if vaccines based on Wuhan-Hu-1 S alone will provide satisfactory protection, either as boosters in experienced individuals or as an initial vaccine in naive individuals (mainly children). The recent evaluation of intranasal vaccine administration could also be important to not only prevent severe disease but also curtail viral infection and transmission through induction of mucosal immunity (*59, 60*). For these reasons, it is important to monitor new variants, assess the effectiveness of currently available vaccines, and continue to test and implement new vaccination strategies that may provide stronger, longer lasting, or broader protection against SARS-CoV-2 and the entire sarbecovirus subgenus (*38, 61, 62*).

## Author contributions

J.E.B. and D.V. conceived the project and designed experiments. J.E.B. performed all experiments. J.E.B. and K.R.S. produced pseudotyped viruses. A.C.W., I.G.K., J.K.L, N.M.F., K.A., A.S., C.M.P., J.D., Z.Z., D.W., A.S, S.C., N.T.I., D.C., J.G., R.G., and H.Y.C. provided unique reagents. D.V. supervised the project and obtained funding. J.E.B. and D.V. analyzed the data and wrote the manuscript with input from all authors.

## Acknowledgements

We thank Hideki Tani (University of Toyama) for providing the reagents necessary for preparing VSV pseudotyped viruses. This study was supported by the National Institute of Allergy and Infectious Diseases (DP1AI158186 and HHSN272201700059C to D.V.), the National Institute of General Medical Sciences (R01GM120553 to D.V.), a Pew Biomedical Scholars Award (D.V.), an Investigators in the Pathogenesis of Infectious Disease Awards from the Burroughs Wellcome Fund (D.V.), Fast Grants (D.V.), the Bill & Melinda Gates Foundation (OPP1156262 to D.V.), the University of Washington Arnold and Mabel Beckman cryoEM center and the National Institute of Health grant S10OD032290 (to D.V.) and grant U01 AI151698 for the United World Antiviral Research Network (UWARN) as part of the Centers for Research in Emerging Infectious Diseases (CREID) Network. Additionally, work was supported by NIH NIAID grant CCHI AI142742 (S.C., A.S), NIAID Contract No. 75N9301900065 (A.S, D.W), and U01 CA260541-01 (D.W). D.V. is an Investigator of the Howard Hughes Medical Institute.

## Supplementary Material

**Table S1.** SARS-CoV-2 BA.1 and BA.2 spike mutations as compared to Wuhan-Hu-1.

| BA.1 | Both | BA.2 |
| --- | --- | --- |
| A67V, del69/70, T95I,<br>del143/145, N211I,<br>del212/212, S371L, G496S,<br>Q498R, T547K, N856K,<br>L981F | G142D, G339D, S373P,<br>S375F, S477N, T478K,<br>E484A, Q493R, Q498R,<br>N501Y, Y505H, D614G,<br>H655Y, N679K, P681H,<br>N764K, D796Y, Q954H,<br>N969K | T19I, L24S, del25/27, V213G,<br>S371F, T376A, D405N,<br>R408S, K417N, N440K |

**Table S2.**
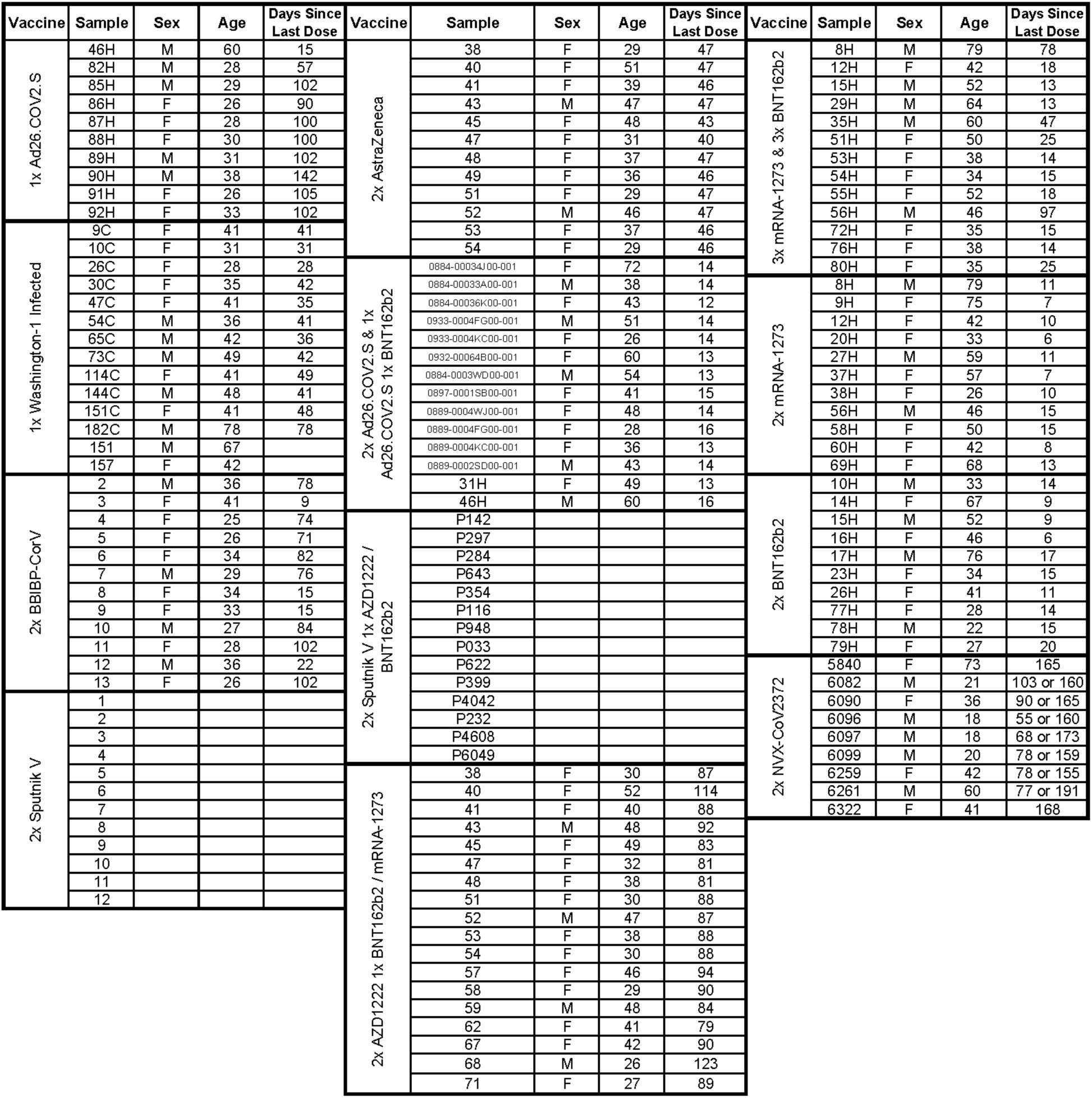
Demographics data of enrolled plasma donors.

**Fig. S1.**
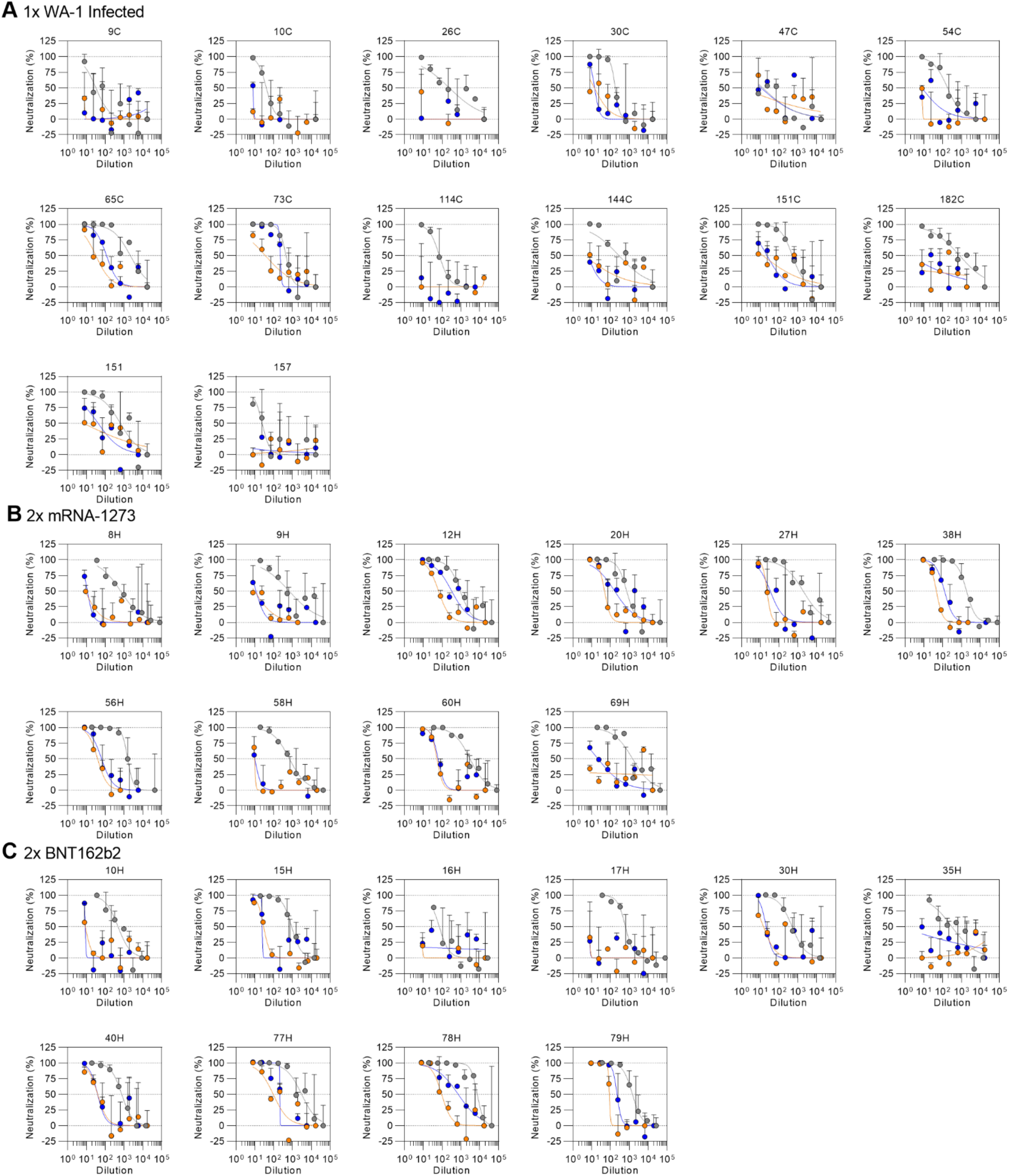
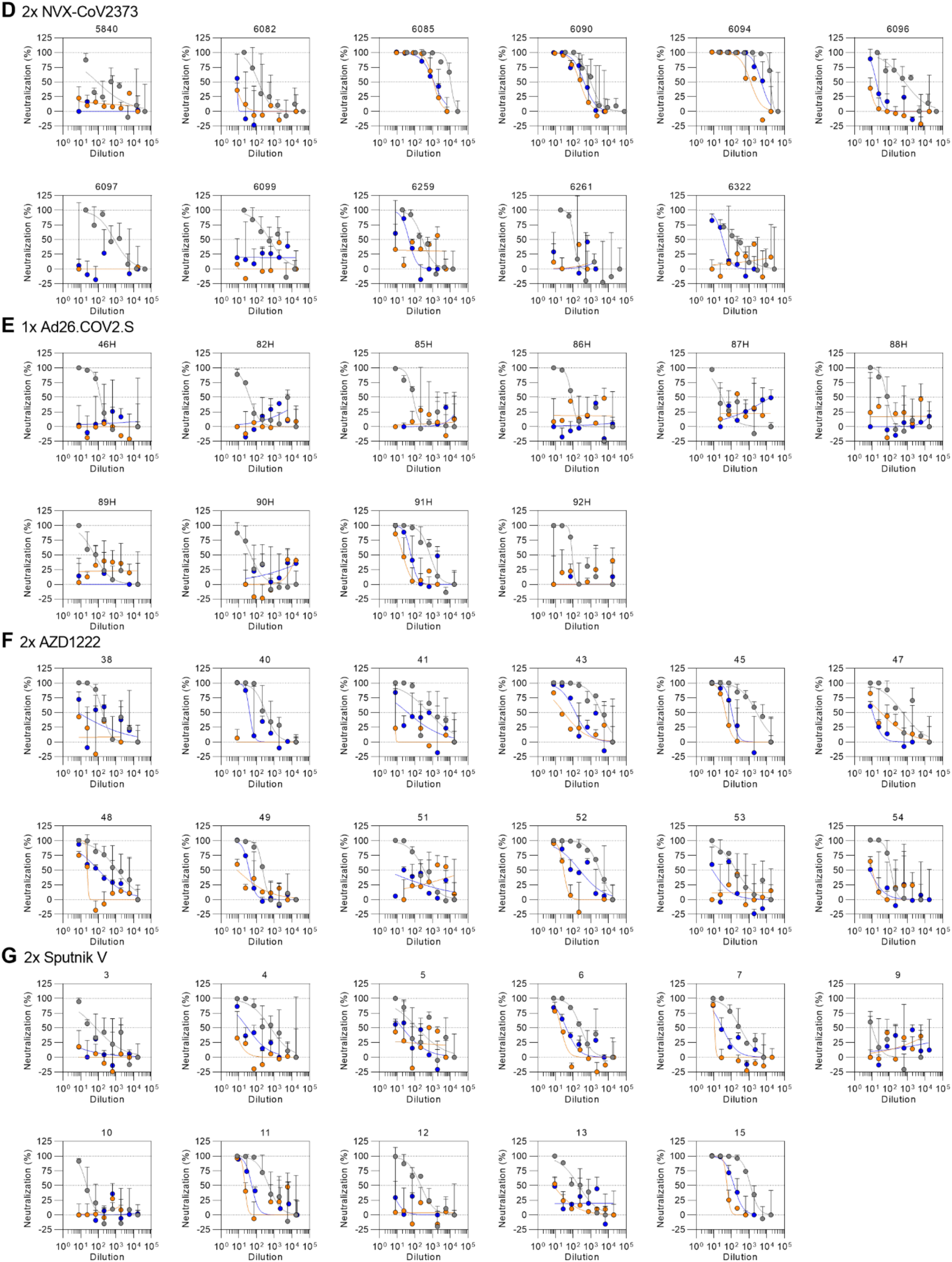
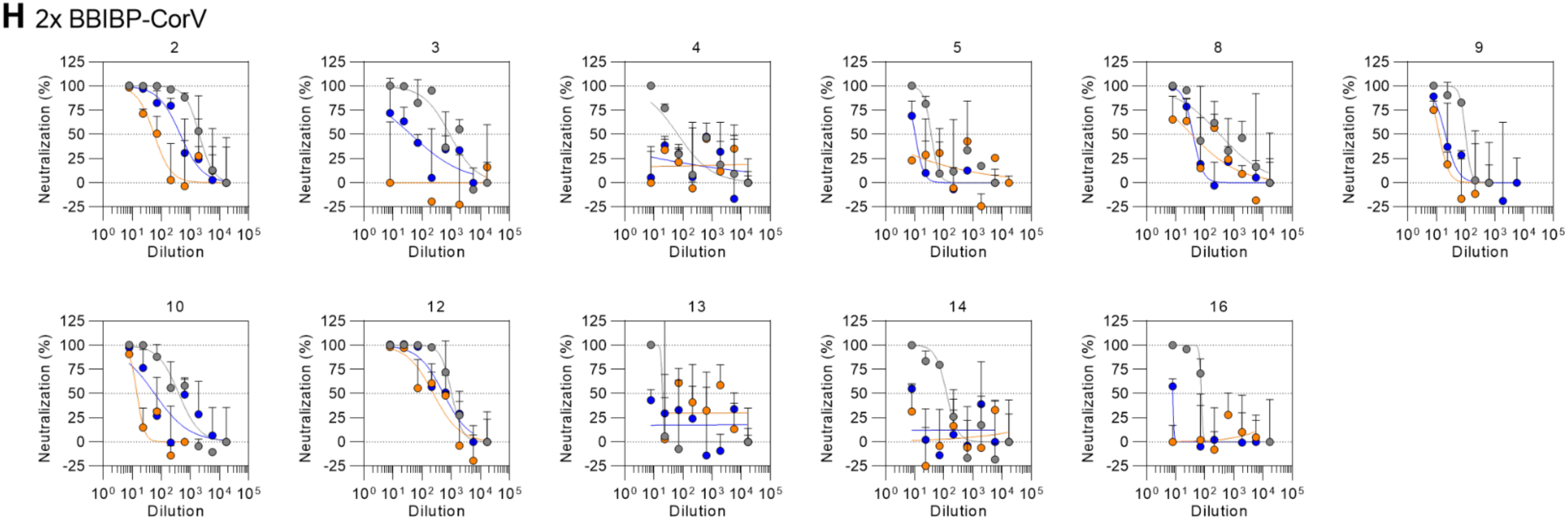
Normalized neutralization curves using VSV pseudovirus containing the SARS-CoV-2 G614 spike (gray), BA.1 spike (orange), or BA.2 spike (blue) on VeroE6-TMPRSS2 cells using plasma from subjects after their primary vaccine series.

**Fig. S2.**
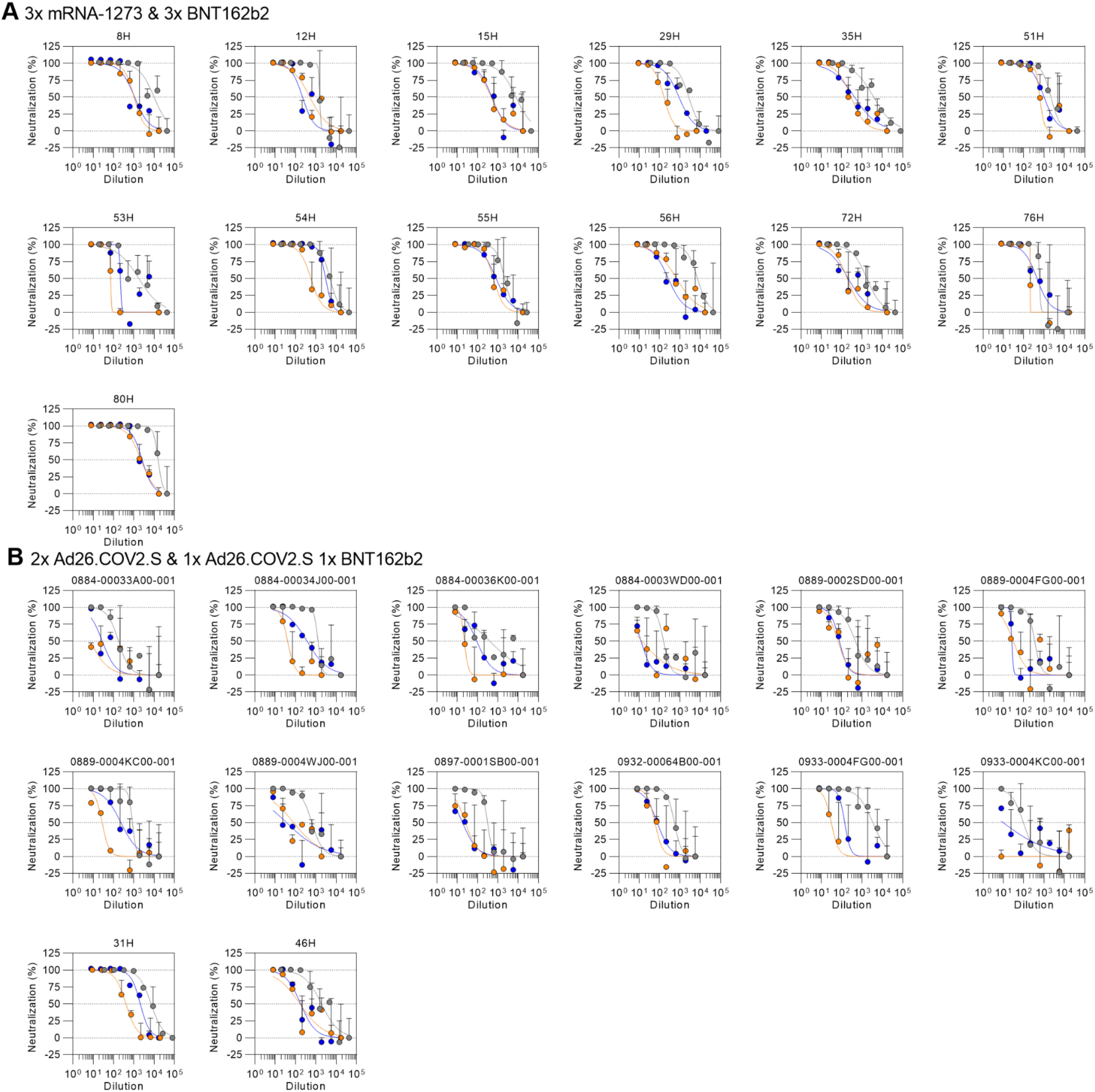
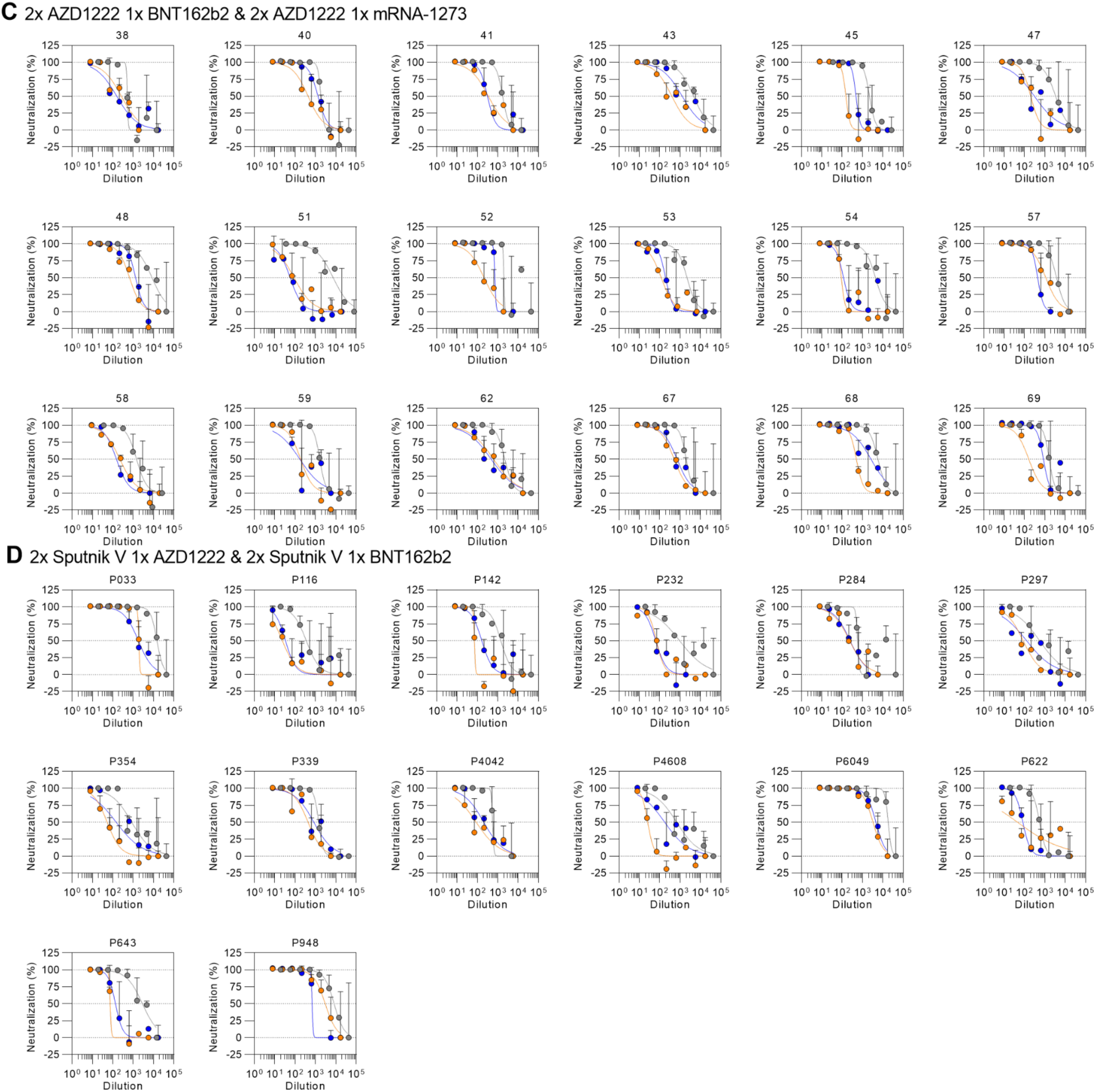
Normalized neutralization curves using VSV pseudovirus containing the SARS-CoV-2 G614G (gray), BA.1 spike (orange), or BA.2 spike (blue) on VeroE6-TMPRSS2 cells using plasma from subjects after receiving a booster dose.

## Methods

### Cell lines

Cell lines used in this study were obtained from ThermoFisher Scientific (HEK293T) or were kindly gifted by Florian Lempp (Vero-TMPRSS2 cells(*40*)). None of the cell lines used were authenticated or tested for mycoplasma contamination.

### Sample donors

Convalescent plasma, Ad26.COV2.S, and some BNT162b2 samples were obtained from the HAARVI study approved by the University of Washington Human Subjects Division Institutional Review Board (STUDY00000959). mRNA-1273 and BNT162b2 samples were obtained from individuals enrolled in the UWARN: COVID-19 in WA study approved by the University of Washington Human Subjects Division Institutional Review Board (STUDY00010350). AZD1222 samples were obtained from the PolImmune-COVID study conducted by INGM and IRCCS Ca’ Granda Ospedale Maggiore Policlinico of Milan, approved by INMI “Lazzaro Spallanzani” Ethics Committee (286_2021). Samples from NVX-CoV2373 immunized individuals were collected in the San Diego region by the La Jolla Institute for Immunology. This work was approved by the institutional review boards (IRB) of the La Jolla Institute (IRB#: VD-214). Sputnik V samples were obtained from healthcare workers at the hospital de Clínicas “José de San Martín”, Buenos Aires, Argentina. BBIBP-CorV samples were obtained from Aga Khan University, Karachi, Pakistan. Demographic data for these individuals are summarized in Table S2.

### Plasmid construction

SARS-CoV-2 G614 S (YP 009724390.1) S gene was placed into the HDM vector with a 21 residue C-terminal deletion, as previously described (*3, 38, 63*). The plasmids encoding the SARS-CoV-2 Omicron S variants BA.1 and BA.2 were generated by overlap PCR mutagenesis of the wildtype plasmid, pcDNA3.1(+)-spike-D19 (**Table S1**) (*64*).

### Pseudotyped VSV production

SARS-CoV-2 G614 and Omicron BA.1 and BA.2 pseudotypes were prepared as previously described (*3*). Briefly, HEK-293T cells seeded in poly-D-lysine coated 100 mm dishes at ∼75 % confluency were washed five times with Opti-MEM and transfected using 24 μg of the S glycoprotein plasmid with Lipofectamine 2000 (Life Technologies). After 5 h at 37°C, media supplemented with 20% FBS and 2% PenStrep was added. After 20 hours, cells were washed five times with DMEM and cells were transduced with VSVΔG-luc before a 2 h incubation at 37°C. Infected cells were then washed an additional five times with DMEM prior to adding media supplemented with anti-VSV-G antibody (I1-mouse hybridoma supernatant diluted 1:25, from CRL-2700, ATCC) to reduce parental background. After 18-24 h, the supernatant was harvested and clarified by low-speed centrifugation at 2,500 g for 10 min. The supernatant was then filtered (0.45 μm) and concentrated 10 times using a 30 kDa centrifugal concentrator (Amicon Ultra). The pseudotypes were then aliquoted and frozen at −80 °C.

### Pseudotyped VSV neutralization assay

To evaluate neutralization of SARS-CoV-2 G614 and Omicron BA.1 and BA.2 pseudotypes by plasma of vaccinees or previously infected individuals, Vero-TMPRSS2 cells in DMEM supplemented with 10% FBS, 1% PenStrep, and 8 ug/mL puromycin were seeded at 60-70% confluency into white clear-bottom 96 well plates (Corning) and incubated at 37°C. The following day, a half-area 96-well plate (Greiner) was prepared with eight 3-fold serial plasma dilutions. An equal volume of DMEM with 1:25 pseudovirus and 1:25 anti-VSV-G antibody (I1-mouse hybridoma supernatant from CRL-2700, ATCC) was then added to the half-area plate. The mixture was incubated at room temperature for 20-30 minutes. Media was removed from the cells and 40 μL from each well (containing plasma and pseudovirus) was transferred to the 96-well plate seeded with Vero-TMPRESS2 cells and incubated at 37°C for 2 h. After 2 h, an additional 40 μL of DMEM supplemented with 20% FBS and 2% PenStrep was added to the cells. After 16-20 h, 40 μL of One-Glo-EX substrate (Promega) was added to each well and incubated on a plate shaker in the dark for 5 min. Relative luciferase units were read using a Biotek plate reader. Relative luciferase units were plotted and normalized in Prism (GraphPad): 100% neutralization being cells lacking pseudovirus and 0% neutralizing being cells containing virus but lacking plasma. Prism (GraphPad) nonlinear regression with “[inhibitor] versus normalized response with a variable slope” was used to determine ID50 values from curve fits with 2-3 repeats. 2-4 biological replicates were carried out for each sample.

